# On the move: Redox –dependent protein relocation

**DOI:** 10.1101/588269

**Authors:** Christine H. Foyer, Alison Baker, Megan Wright, Imogen A. Sparkes, Amna Mhamdi, Jos H.M. Schippers, Frank Van Breusegem

## Abstract

Compartmentation of proteins and processes is a defining feature of eukaryotic cells. The growth and development of organisms is critically dependent on the accurate sorting of proteins within cells. The mechanisms by which cytosol-synthesized proteins are delivered to the membranes and membrane compartments have been extensively characterised. However, the protein complement of any given compartment is not precisely fixed and some proteins can move between compartments in response to metabolic or environmental triggers. The mechanisms and processes that mediate such relocation events are largely uncharacterized. Many proteins can in addition perform multiple functions, catalyzing alternative reactions or performing structural, non-enzymatic functions. These alternative functions can be equally important functions in each cellular compartment. Such proteins are generally not dual targeted proteins in the classic sense of having targeting sequences that direct *de novo* synthesised proteins to specific cellular locations. Accumulating evidence suggests that redox post-translational modifications (PTMs) can control the compartmentation of many such proteins, including antioxidant and/or redox associated enzymes.

## INTRODUCTION

Many proteins perform multiple unrelated functions, often in different locations. These are often referred to as moonlighting proteins. Some of these proteins have been known for decades. However, the number and diversity of proteins that either can have different functions in the same intracellular compartment or that can move from one compartment to another to fulfil different functions has increased enormously in recent years, aided by development and application of bioinformatic (Chapple *et al.*, 2015) proteomic (Thul *et al.*, 2017) and cell imaging techniques (Chong *et al.*, 2015; Thul *et al.*, 2017). These reveal the extent to which proteins can show multiple subcellular localisations (up to 50% of cellular proteins), as well as how these may change in response to cellular perturbation (Chong *et al.*, 2015), including in disease states like cancer in animals (Min *et al.*, 2016) and stress responses in plants (Sun et al. 2018). Several metabolic enzymes are known to move into the nucleus affecting epigenetic modifications (Boukouris *et al.*, 2016) and histone expression (He *et al.*, 2013) providing a link between metabolism and gene expression. Whilst most studies have been conducted on yeast and mammalian cells, evidence for proteins with multiple, largely unrelated functions in plants is also incontrovertible (**Table I**). A recent study in Arabidopsis identified a number of metabolic enzymes as members of the RNA binding protein repertoire (Marondedze *et al.*, 2016). Not all proteins that move moonlight, and not all proteins that moonlight move compartments to do so. For example, L-galactono-1, 4-lactone dehydrogenase has dual functions in plant mitochondria. Firstly, as an enzyme it is responsible for the synthesis of ascorbic acid, and as a chaperone it is essential for the assembly of respiratory complex I (Schimmeyer *et al.*, 2016). Similarly plastid NAD dependent malate dehydrogenase has a non-enzymatic function stabilising the FtsH12 component of the inner envelope AAA ATPase (Schreier *et al.*, 2018).

Regulated protein relocation provides a robust and flexible mechanism for metabolic, genetic and epigenetic regulation in response to metabolic stimuli and environmental cues. Such responses often entail shifts in cellular redox homeostasis that lead to both oxidative and reductive events that shift protein functions and compartmentation. One paradigm for such redox-related changes is NPR1, which is a master regulator of salicylic acid (SA)-mediated systemic acquired resistance (SAR) leading to broad-spectrum disease resistance in plants (Mou *et al.*, 2003). A second paradigm is organelle-to-nucleus retrograde signalling pathways that allow cells to adapt to changes in metabolic state, often in a redox-dependent manner (Boukouris *et al.*, 2016; Monaghan and Whitmarsh, 2015). Redox cues and associated PTMs are often fundamental regulators of alternative protein functions and localization. However, the extent of this phenomenon, what makes proteins move and the mechanisms by which they do so remains largely obscure.

## MECHANISMS OF PROTEIN MOVEMENT IN PLANTS

### ROS triggered post translational modifications (PTMS)

All cells use oxidative catabolic processes to release energy and anabolic reductive process to assimilate energy in synthetic reactions. Redox signalling was one of the first regulatory pathways to evolve to avoid “boom” and “bust” scenarios in energy availability and usage (Foyer and Allen, 2003). Examples of oxidative stress triggered protein movement are known in yeast (e.g. Superoxide Dismutase SOD1 (Tsang *et al.*, 2014)) and mammalian cells (e.g NRF2, CLK-1, glyceraldehyde 3-phosphate dehydrogenase (GAPDH) reviewed in (Min *et al.*, 2016; Monaghan and Whitmarsh, 2015) and results in these proteins exhibiting moonlighting activity within the nucleus.

Reactive Oxygen Species (ROS) serve as key regulators of a diverse range of important functions in plants. In particular, they serve as signalling molecules that control plant growth through processes such a mitosis, cell expansion and differentiation (Mhamdi and Van Breusegem, 2018). ROS are essential signals produced by cellular energy metabolism and by specific enzymes such as NADPH oxidases in order to modulate redox-sensitive processes (Schmidt and Schippers, 2015). Chloroplasts, mitochondria and peroxisomes are thus not only the essential sites of metabolic energy production and utilization, but also important sources of ROS and other redox regulators that influence nearly every aspect of cell biology (Noctor and Foyer, 2016). While cells regulate redox processes in a compartment-specific manner, redox PTMs may also be used to regulate the movement of proteins between compartments. Moreover, proteins such as peroxiredoxins (PRX) that readily undergo redox PTMS in their roles as ROS scavengers are moonlighting enzymes that have evolved to support multiple functions (acting as peroxidases, signalling proteins and chaperones) under optimal and stress conditions. Like other redox proteins, whose functions are supported by thiol-based biochemistry, PRX can interact with multiple cellular partners in animals and plants, from thioredoxins to transcription factors (Liebthal *et al.*, 2018). They are thus a paradigm for moonlighting proteins

Assessing the protein-protein interactions that are involved in the functions of PRXs and other redox regulated proteins such as NPR1 entails considerations of the interdependent facets of redox state and oligomeric structure. Redox PTMs on protein cysteines are formed non-enzymatically via promiscuous reactive species, including ROS, reactive nitrogen species (RNS), and other radicals or electrophilic lipids. There is growing appreciation that redox PTMs are site-specific, governed by the microenvironment of cysteine residues, and subject to temporal and spatial control. Small molecule and protein-based fluorescent sensors have shown that eukaryotic cells tightly control the location of reactive species, proteins and redox state across compartments (Kaludercic *et al.*, 2014), and that this balance is, for example, altered during ageing in the model organism *C. elegans* (Kirstein *et al.*, 2015). Recent evidence suggests that ROS and redox cues modify microtubule orientation and behaviour within plant cells (Dang *et al.*, 2018), as well as the operation of protein import machineries (reviewed in (Bolter *et al.*, 2015; Ling and Jarvis, 2015). Redox PTMs not only control activities and binding partners but also the compartmentation of many proteins, including antioxidant and/or redox associated enzymes (**Box I**), as discussed in detail below.

### Protein Import and Export

The molecular mechanisms of protein import into mitochondria, chloroplasts and peroxisomes have now been established and the importance of the accuracy of these processes underscored by the realization that defects result in human disease. Recent work has revealed that protein import can be regulated at several levels; from modification of individual precursor proteins to prevent or alter their targeting, to regulated interaction with binding partners, and modification of the import apparatus by phosphorylation or ubiquitination to alter its activity (Bolter *et al.*, 2015; Harbauer *et al.*, 2014; Ling *et al.*, 2012) (**Figure 1**). Such processes allow the location of proteins to change in response to changes in cellular state. For example, in *C. elegans* the transcription factor ATF1 is imported into mitochondria and degraded by a Lon protease but, when import is decreased, ATF1 relocates to the nucleus and induces an unfolded protein response (Nargund *et al.*, 2012). In mammals, import of the protein catalase into the peroxisome is redox regulated and under stress conditions the peroxisome import receptor PEX5, retains catalase in the cytosol (Walton *et al.*, 2017). PEX5 cycling between peroxisome and cytosol is regulated by ubiquitination of a conserved Cys, and in mammalian cells reduced glutathione can deubiquitinate the receptor (Grou *et al.*, 2009). Intriguingly, an old observation that NADPH but not NADH inhibits protein import hints at the importance of redox balance for protein import into plant peroxisomes as well (Pool *et al.*, 1998). Retrograde signalling from organelles to the nucleus to integrate cellular activities is well established, and modulation of chloroplast import activity is important in response to biotic and abiotic stress (de Torres Zabala *et al.*, 2015; Ling and Jarvis, 2015).

**Figure 1.**
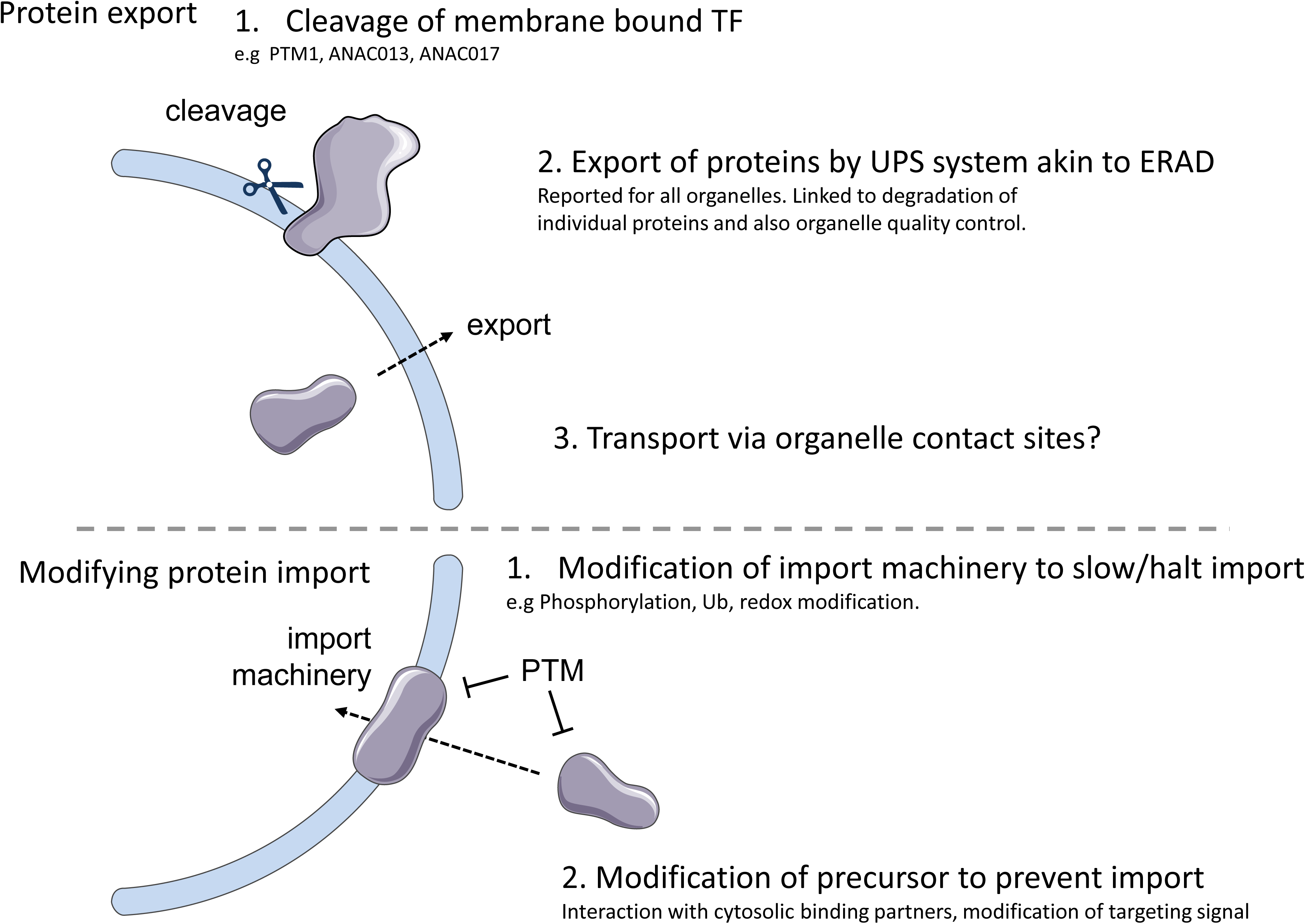
Potential mechanisms of protein relocation. Proteins can potentially change their cellular localisation by a number of mechanisms. Proteins which have been inserted into an organelle membrane can be released by regulated proteolysis as described for the chloroplast envelope localised PTM1 and ER membrane localised ANAC013 and ANAC017. All organelles appear to have an ER associated degradation (ERAD)-like pathway which exports proteins in an ubiquitin dependent manner for degradation by the proteasome. Whether proteins can be exported and escape degradation to be retargeted elsewhere in the cell is currently unknown. Ubquitination on membrane components can also lead to organelle turnover. Transport by direct organelle contacts is also a possible mechanism. Proteins normally targeted to an organelle can be prevented from import through either modification of the import machinery or modification of the cyctosolic precursor form of the protein. This can include post translational modifications (PTMs) which can modify the targeting signal or affect interactions with other binding partners. See text for further details.

As well as regulating the import of proteins, it has become apparent that proteins can be exported from mitochondria, chloroplasts and peroxisomes as well as the endoplasmic reticulum (ER) (**Figure 1**). Such export not only drives degradation of organellar proteins via the cytosolic Ubiquitin-proteasome system (UPS), but also provides a link to organelle quality control (Bragoszewski *et al.*, 2017; Kao *et al.*, 2018; Ling and Jarvis, 2016). While the release of transcription factors from cellular membranes by regulated proteolysis is a well-known response to stress in both animals and plants (Seo *et al.*, 2008; Sun *et al.*, 2011), potentially, protein export and retargeting could also provide a means of signalling and genetic regulation. To date this has only been proposed/described for a handful of proteins and the mechanism(s) by which this occurs and is regulated are still obscure (Foyer *et al.*, 2014).

### Candidates as a paradigm for redox regulated movement in plants

#### NPR1 and ROXY proteins

NPR1 shares structural and functional characteristics with mammalian immune co-factor I k B and the transcription factor NF-k B, suggesting cross-kingdom conservation of central immune responses (Sun *et al.*, 2018). NPR1 is a master regulator of SA perception. Relocation of NPR1 to the nucleus is essential for its function in regulating *PR* gene expression. NPR1 resides in a large disulfide-bonded oligomeric complex in the cytoplasm in the absence of stress. However, SA accumulation leads to reduction of the intermolecular disulfide bonds by thioredoxin (TRX; (Tada *et al.*, 2008), releasing monomeric NPR1. Reduced NPR1 is then imported into the nucleus (Mou *et al.*, 2003). This process involves phosphorylation at serine 589 (S589) by SnRK2.8, which is important for NPR1 nuclear localization. After entering the nucleus, phosphorylation of NPR1 at serine 55 and serine 59 (S55/59) promotes its association with transcription factors such as WRKY and TGA in a redox-dependant manners leading to the expression of pathogenesis-related (PR) genes. Similarly, the glutaredoxin ROXY1 and its homologue ROXY2 are found in the nucleus and cytoplasm (Delorme-Hinoux *et al.*, 2016). In the nucleus, ROXY1 plays a key role in petal development interacting with TOPLESS in a redox-dependent manner, and with TGA2, TGA3, TGA7 and PERIANTHIA. However, in the case of ROXY1 there is no evidence as yet of redox-regulated movement between the nucleus and cytoplasm.

**GAPDH** is considered to be a quintessential example of a moonlighting protein (Sirover, 2012, 2014). It has multiple functions in animals such as DNA stability and control of gene expression, autophagy and apoptosis, in addition to its classic role in glycolysis. The functions of GAPDH in the plant nucleus are not clear, but it may act as a coactivator for gene expression (Hildebrandt *et al.*, 2015). Redox PTMs to the cytosolic GAPDH protein in animals, which block enzyme activity, promote novel cell signalling and transcription functions in the nucleus (Yang and Zhai, 2017) (Zaffagnini *et al.*, 2013). Several GAPDH isoforms exist in different subcellular localizations in plants (Holtgrefe *et al.*, 2008). In particular, the activity and localization of the cytosolic GAPDH isoform (GapC) is controlled by cellular redox state (Bedhomme *et al.*, 2012). Since GapC is also localized in the nucleus, it is suggested that redox modification facilitates transfer to the nucleus in plants as it does in animals (Ortiz-Ortiz *et al.*, 2010). However, the mechanism of nuclear translocation of GapC is unknown although it is thought to involve *S*-sulfhydration, a process that reversibly regulates the function of this protein, in a manner similar to that described in mammalian systems (Aroca *et al.*, 2015). However, GapC undergoes *S*-nitrosylation, *S*-glutathionylation, *S*-sulfhydration, *S*-sulfenylation as well as other modifications that all occur on the same cysteine residue (Aroca *et al.*, 2017; Bedhomme *et al.*, 2012; Lindermayr *et al.*, 2005; Waszczak *et al.*, 2014). Thus, how each type of PTM modifies GapC to shrift location and/or alternative instigate non-metabolic functions remains to be determined.

#### Catalase

(CAT) is a peroxisomal enzyme whose import in mammals is redox-regulated (Walton *et al.*, 2017) and in yeast is dependent on carbon source (Horiguchi *et al.*, 2001). In plants it is classically known as a <u>peroxisomal</u> enzyme but recent evidence suggests that the compartmentation of this central antioxidant enzyme may be more dynamic than the literature acknowledges. The role of CAT as a central ‘redox guardian’ is well established (Mhamdi *et al.*, 2012). Plant catalases have been shown to interact with a variety of <u>cytosolic</u> proteins including calmodulin (Yang and Poovaiah, 2002), calcium-dependent protein kinase 8 (CDPK8) (Zou *et al.*, 2015), salt overly sensitive 2 (SOS2) (Verslues *et al.*, 2007), lesion stimulating disease1 (LSD1) (Li *et al.*, 2013), receptor like cytoplasmic kinase STRK1 (Zhou *et al.*, 2018) and no catalase activity 1 (NCA1) (Hackenberg *et al.*, 2013; Li *et al.*, 2015) (**Figure 2**). All are integral stress signalling proteins. The *nca1* mutants, which lack a functional CAT, are hypersensitive to abiotic stresses. Similarly, the *cat2* mutant of Arabidopsis, which lacks the predominant leaf isoform that is essential for the metabolism of H_2_O_2_ produced by photorespiration, activates a wide range of salicylic acid (SA) and jasmonic acid (JA)-dependent responses and displays day-length dependent localised programed cell death (PCD) and resistance to pathogens (Queval *et al.*, 2010). CAT can also be a target for pathogen encoded-effector proteins (Mathioudakis *et al.*, 2013; Murota *et al.*, 2017). The fungal effectors PsCRN115 and PsCRN63 both traffic CAT to the nucleus but have opposite biochemical and physiological effects. PsCRN115 stabilises catalase, decreases H_2_O_2_ and reduces PCD, whereas PsCRN63 destabilises catalase increases H_2_O_2_ and increases PCD (Zhang *et al.*, 2015). We consider that the CAT interactome with different stress signalling and PCD proteins provides a paradigm for the study for protein relocation. We propose that the location of cytosolically-synthesised CAT is determined by competition among different potential-binding partners as a consequence of reduced import into peroxisomes and/or increased retention of CAT in the cytosol. While sensitivity of peroxisomal protein import to redox status is likely to impact import of all peroxisome proteins, CATALASE which has a non-canonical targeting signal (Mhamdi *et al.*, 2012) (Rymer *et al.*, 2018) may be more sensitive and indeed PEX5, the major peroxisome import receptor, has been proposed to specifically retain mammalian catalase in the cytosol under conditions of oxidative stress (Walton *et al.*, 2017). This property, combined with the potential to interact with an array of cytosolic proteins as shown in **Figure 2** could allow swift control of catalase localisation between compartments in such a way as to influence various redox signalling pathways.

**Figure 2.**
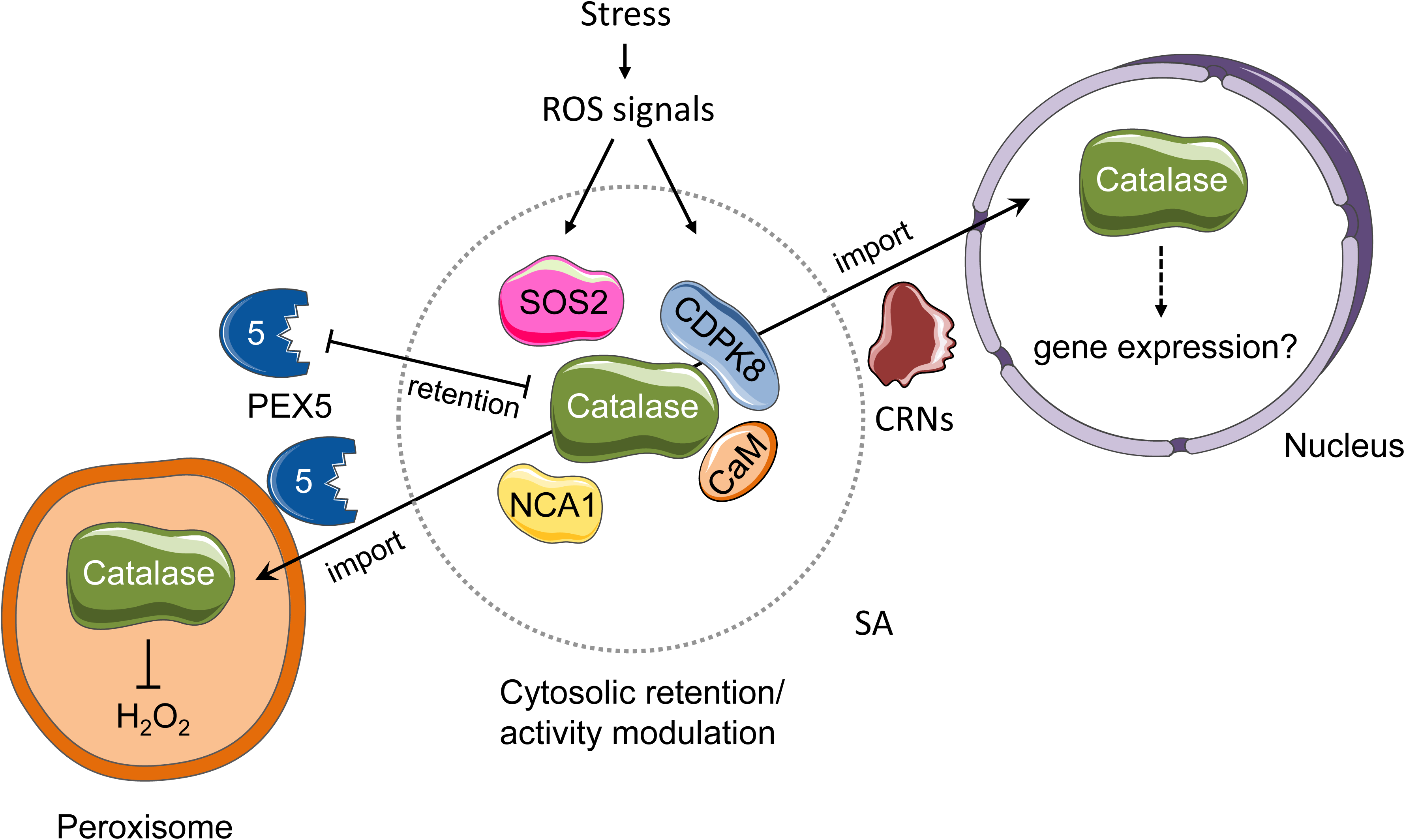
Switching partners: model for regulation of catalase localisation through interaction with different binding proteins. Several cytosolic proteins have been reported to interact with plant catalases. Redox mediated PTMs could alter the affinity of catalase for different binding partners leading to a change in distribution between peroxisomes, cytosol and nucleus. See text for further details.

#### WHIRLY1

(WHY1) is a member of a small family of ssDNA binding proteins that are specific to the plant kingdom (Desveaux *et al.*, 2005; Desveaux *et al.*, 2004). The nuclear-encoded WHY1 protein is targeted to chloroplasts and the nucleus, the nuclear form having the same molecular mass as the processed chloroplast form. In the chloroplasts, WHY1 binds to both DNA and RNA and regulates chloroplast development, plastome copy number and is required for plastome gene expression, intron splicing, ribosome formation and chloroplast to nucleus signaling (Comadira et al., 2015; Prikryl *et al.*, 2008). In the nucleus, WHY1 functions in the transcription of senescence and defence genes as well as in the maintenance of telomeres (Yoo *et al.*, 2007). The partitioning of WHY1 between the chloroplasts and nucleus changes during leaf development, WHY1 being predominantly in the chloroplasts of young leaves, while in senescing leaves the protein is localized mainly in the nucleus (Ren *et al.*, 2017). This partitioning is regulated at least in part by phosphorylated of WHY1 in the cytosol by a serine/threonine SNF1-related protein kinase called calcineurin B-Like-Interacting Protein Kinase14 (CIPK14). Phosphorylation of WHY1 results in transport to the nucleus (Ren *et al.*, 2017). However, other studies using epitope tagged, transplastomically expressed WHY1 have reported that WHY1 can move from the chloroplasts in the nuclei (Isemer *et al.*, 2012). WHY1 may therefore move from plastids to nucleus upon redox signals (Foyer *et al.*, 2014).

#### Membrane bound transcription factors

As well as soluble proteins, membrane located proteins can be cleaved from their membrane anchor in response to an appropriate signal and relocated (**Figure 1**). Often these proteins function as transcription factors once liberated from the membrane. *ANAC013* and *ANAC017* encode Arabidopsis transcription factors belonging to the *NON APICAL MERISTEM/ARABIDOPSIS TRANCRIPTION ACTIVATION FACTOR/CUP SHAPED COTYLEDON* (*NAC*) family. These transcription factors mediate ROS-related retrograde signalling originating from mitochondrial complex III. Both proteins contain putative transmembrane domains. They were identified via one hybrid assays as binding to a conserved cis acting regulatory sequence the MDM which mediates mitochondrial retrograde regulation (MRR) during oxidative stress (De Clercq *et al.*, 2013). *ANAC013* was shown via chromatin immunoprecipitation to transactivate MDM in planta and upregulate several *MITOCHONDRIAL DYSFUNCTION STIMULATION* (*MDS*) genes. GFP-ANAC013 was partially processed and nuclear localised, but while difficult to detect there was a suggestion that the full length protein is ER targeted (De Clercq *et al.*, 2013). *ANAC017* was also identified in a screen for loss of response to mitochondrial dysfunction (Ng *et al.*, 2013). It is targeted to the ER and dual tagging experiments showed it is cleaved upon antimycin A treatment, the N terminal part locates to the nucleus whilst the C terminal part remained ER associated. *ANAC017* function was essential for hydrogen peroxide mediated stress signalling (Ng *et al.*, 2013). Upon perception of redox signals, *ANAC013* and *ANAC017* are released from the ER and translocated to the nucleus, where they activate MDS genes such as alternative oxidases (*AOXs*), *SOT12*, and *ANAC013*. The latter provides positive feedback regulation of the signalling pathway with enhancement of the signal. The ROS-dependent signalling pathways from chloroplasts and mitochondria merge at RADICAL-INDUCED CELL DEATH1 (RCD1), a nuclear protein that suppresses the activities of the ANAC013 and ANAC017 transcription factors (Shapiguzov *et al.*, 2019).

The chloroplast bound plant homeodomain transcription factor PTM plays a crucial role in chloroplast signalling to the nucleus. Mutants defective in this gene show aberrant responses to treatments such as Norfluazon, high light dibromothymoquonine and Rose Bengal that affect different ROS and the level of reduction of the plastoquinione pool. Full length PTM is located to the chloroplast outer envelope where as a truncated form lacking the TM domain was nuclear. Treatments such as high light and Norfluazon resulted in cleavage of PTM and localisation of the N terminal domain to the nucleus. Processed PTM was shown to activate ABI4 transcription (Sun *et al.*, 2011).

PEX2 is a peroxisome membrane protein with a cytosolically exposed RING domain E3 ligase that regulates the recycling and turnover of the PEX5 import receptor through ubiquitination (Burkhart *et al.*, 2014). Interestingly a mutant of Arabidopsis *PEX2* (*ted3)* was recovered as a suppressor of the photomorphogenesis mutant *det1* (Hu *et al.*, 2002). The mechanism of this remains unknown but an artificially expressed RING domain was found in the nucleus where it interacted with the transcription factor HY5 (Desai *et al.*, 2014). Possible mechanisms could be cleavage of the RING domain and relocation to the nucleus, alternative transcription/translation sites or direct movement between peroxisome and nuclear membrane. Since peroxisomes are important nodes in the cell’s antioxidant network and import is under redox control we speculate that PEX2 relocation could represent a potential mechanism for sensing the redox state of peroxisomes and relaying this information to the nucleus.

### Organelle movement and contact as a mechanism of protein movement

Apart from release of proteins from membranes, prevention of import into or promotion of export from organelles, direct transfer of proteins between membrane bound compartments via membrane extensions and contact sites can occur (Pérez-Sancho *et al.*, 2016) (**Figure 3**). The cytoplasm in plant cells is densely packed and mainly constrained by the vacuole and ER to a narrow cortical zone. Protein transfer between organelles requires regulated release and redirection. Redirection through the cytosol may be slow and prevent bulk delivery. Emerging evidence suggests that the physical interaction between organelles is a requirement for the exchange of small molecules, lipids and proteins in plants as well as in mammals and yeast (Cohen *et al.*, 2018). Coordinated re-arrangement of organelle positioning within the cell could provide a mechanism for shuttling moonlighting proteins between compartments. Targeted ‘protected’ delivery from degradation, or potential reversal of the PTM, could be provided through the formation of a micro-environment between organelles that allows for exchanging proteins through a narrow 10-40nm cytoplasmic zone at the membrane contact site interface. Repositioning of organelles could also allow neighbouring organelles to signal to one another to regulate protein exchange. Organelle movement and positioning, which play essential roles in plant responses to light and other metabolic and environmental stimuli, are linked to the cellular redox status.

**Figure 3.**
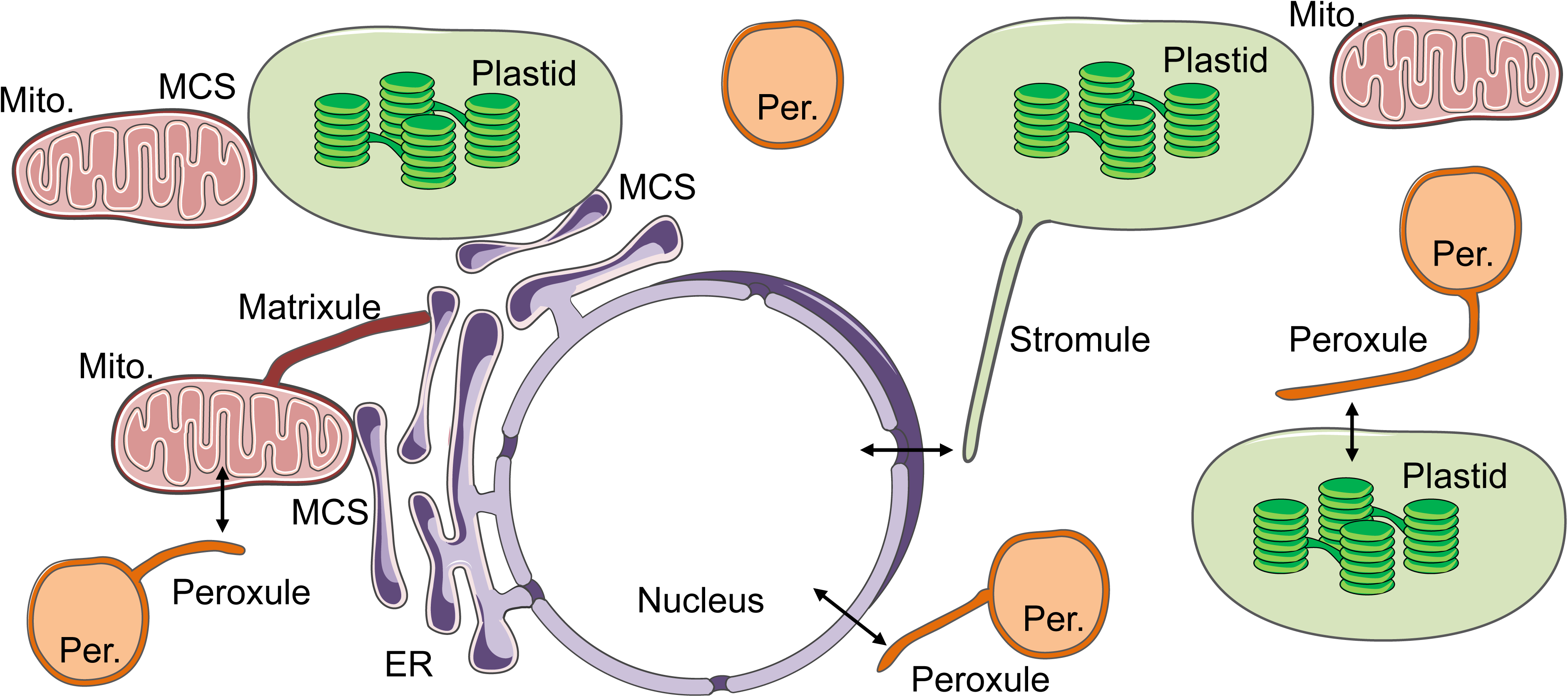
Organelle interactions through protrusions and membrane contact sites. Organelle-organelle interactions in cells occurs by either the formation of membrane contact sites (MCS) between organelles or by the formation of tubular structures by one organelle. MCSs are known to occur between mitochondria and plastids, mitochondria and the ER, and plastids and the ER. In addition, mitochondria, plastids and peroxisomes form tubular structures. In the case of plastids stromules are formed especially in the direction of the nucleus. Mitochondria form matrixules within ER structures and peroxisomes form peroxules in the vicinity of plastids, mitochondria and the nucleus. Both MCS and the tubular structures will mediate communications between organelles by exchanging signaling molecules, metabolites or potentially even proteins.

### Redox-dependent formation of of stromules, matrixules and peroxules

Chloroplasts, mitochondria and peroxisomes are pleomorphic, dynamic organelles that produce tubules upon stress. Like membrane contact sites (MCS) these tubules allow positioning of the organelles in relation to each other within the cell and might be involved in the exchange of metabolites or macromolecues. For example, stroma-filled tubules called stromules **(Figure 4)** extend from the envelope of all plastid types. ROS increase peroxisome speed, resulting in membrane extensions (peroxules), which could facilitate contact with other organelles including chloroplasts (Gao *et al.*, 2016; Rodriguez-Serrano *et al.*, 2016; Rodriguez-Serrano *et al.*, 2009). However, the cargo of the tubular structures and the nature of the potential signals (metabolic or proteinaceous) that are released is largely unknown.

**Figure 4.**
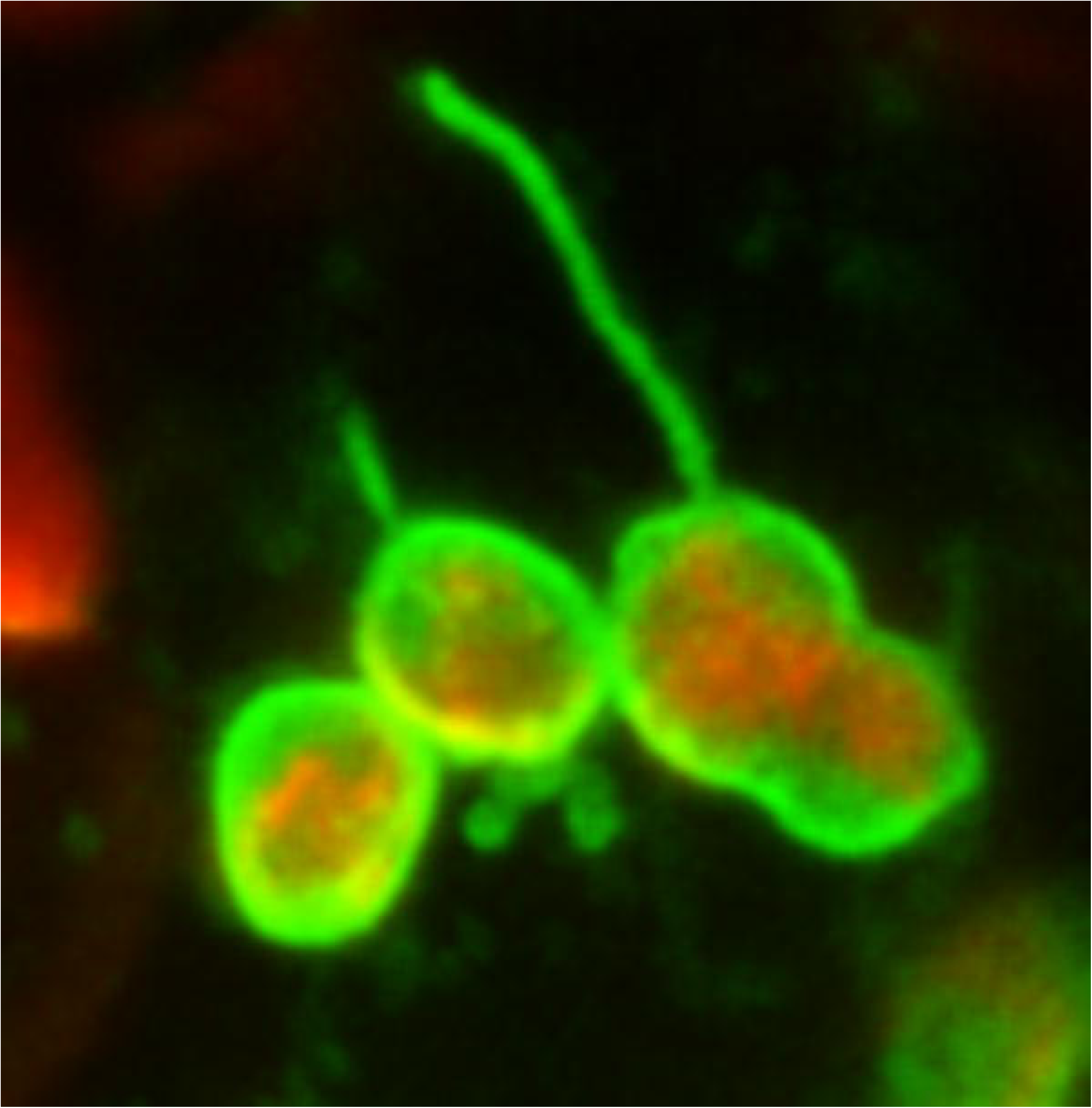
Stromule formation in *N. benthamiana* leaves upon transient over-expression of GFP-tagged plastid outer membrane protein AtLACS9 (At1g77560) *A. tumefaciens* carrying AtLACS9-GFP (Breuers *et al.*, 2012) and a second strain carrying the P19 silencing suppressor construct (Takeda *et al.*, 2002) were co-infiltrated at 0.4 OD each into 7-week-old *N. benthamiana* leaves. Fluorescence imaging was done at 72 hrs. post infiltration with a Zeiss LSM710 confocal microscope. Image is a maximum projection of 10 optical sections. GFP (green); chlorophyll autofluorescence (red).

**Figure 5.**
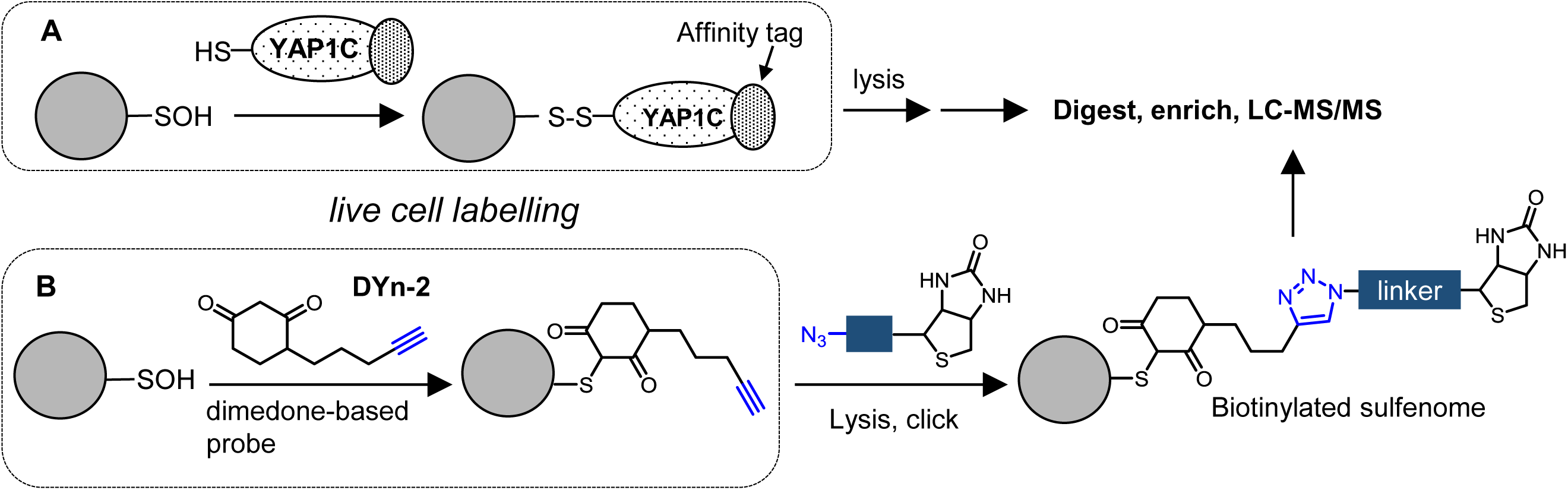
Methods to identify the sulfenome. A. Protein-based probe YAP1C. B. Small molecule-based probe DYn-2. See Box I text for detail.

Stromules allow actin-mediated anchoring of chloroplasts at different locations within the cell to facilitate specific functions. For example, they can extend along microtubules to guide chloroplast movement to the nucleus during innate immunity responses. The application of hydrogen peroxide (H_2_O_2_) resulted in rapid stromule formation in Arabidopsis leaves (Caplan *et al.*, 2015). The accumulation of ROS, like other pro-defense molecules, is sufficient to induce stromule formation leading to the development of direct contact points between the chloroplasts and nuclei (Caplan *et al.*, 2015). In addition, other direct contact sites between chloroplasts and nuclei that are induced by high light have been suggested to allow movement of H_2_O_2_ to the nucleus from attached chloroplasts (Exposito-Rodriguez *et al.*, 2017). Arogenate dehydratase (ADT) 2 which catalyzes the final step in phenylalanine biosynthesis localizes to stromules and also helps in dividing chloroplasts, whilst ADT5 is proposed to traffic to nuclei via stromules, (Bross *et al.*, 2017). Another interesting example of possible organelle to organelle transport of proteins via membrane extensions is the triacyl glycerol lipase SDP1 which is proposed to move from peroxisomes to oil bodies in a tubule- and retromer-dependent process (Thazar-Poulot *et al.*, 2015).

Mitochondria produce structures that are partly homologous to the chloroplast stromules, in (Schmidt *et al.*, 2016) response to light and other stimuli in an endoplasmic reticulum (ER)-mediated manner. The protrusion-driven movement and positioning is considered to promote the inter-compartmental trafficking of metabolites and proteins but there remains a paucity of data on which proteins are trafficked and the mechanisms involved. ROS and redox cues modify microtubule orientation and behaviour within cells, as well as the operation of protein import and export machineries (Schmidt *et al.*, 2016). So far, it remains to be determined if these organelle-derived tubular structures are involved in direct exchange of metabolites or macromolecules between compartments, or might rather have a supportive function in the communication between organelles by acting as an cellular anchor to temporary fix their relative position to each other.

### Conclusions and Perspectives

The regulation of metabolism is shaped by compartmentalization in all cell types. Indeed, the compartmentalization of cell structure is a prerequisite for establishing stable metabolic states of different cell fates (Harrington *et al.*, 2013). Higher plants show a particularly high degree of cellular compartmentalization because of the presence of compartments such as chloroplasts, other plastids and vacuoles. Until recently the paucity of experimental data on subcellular protein distribution has limited our understanding of the capacity and ability of proteins to move between different intracellular compartments. Recent years have seen a step change in our knowledge of proteins that perform more than one cellular function. The term given to such proteins is ‘moonlighting’, but this description is limited because it does not apply to all proteins that move between different cellular compartments. Moreover, it has become increasingly apparent that protein localisation is not fixed and a high proportion of cellular proteins have the potential to move between compartments in response to specific triggers. In some cases this movement is the basis for an alternative cellular function. We have provided a list of examples of proteins that are considered to undergo redox-regulated movements between compartments. This may be the tip of iceberg because there are many proteins in the literature that are suggested to undergo inter-compartmental switching in response to appropriate triggers. Arabidopsis hexokinase 1, for example, which is located at the outer mitochondrial membrane, was suggested to be translocated between mitochondrion and nucleus, upon perception of sugar signals or methyl-jasmonate, in a manner that is linked to mitochondrial ROS production (Claeyssen and Rivoal, 2007; Xiang *et al.*, 2011). At present however we have only a fragmented picture with relatively few well characterised examples of proteins in plants that change compartment in order to moonlight, and the mechanisms by which they do so are largely unexplored. Redox PTMs are likely to be a key driver for inter-compartmental shifts of antioxidant and redox-regulated proteins that can integrate metabolic processes and influence genetic and epigenetic controls of plant growth and stress tolerance.

**Table 1.**
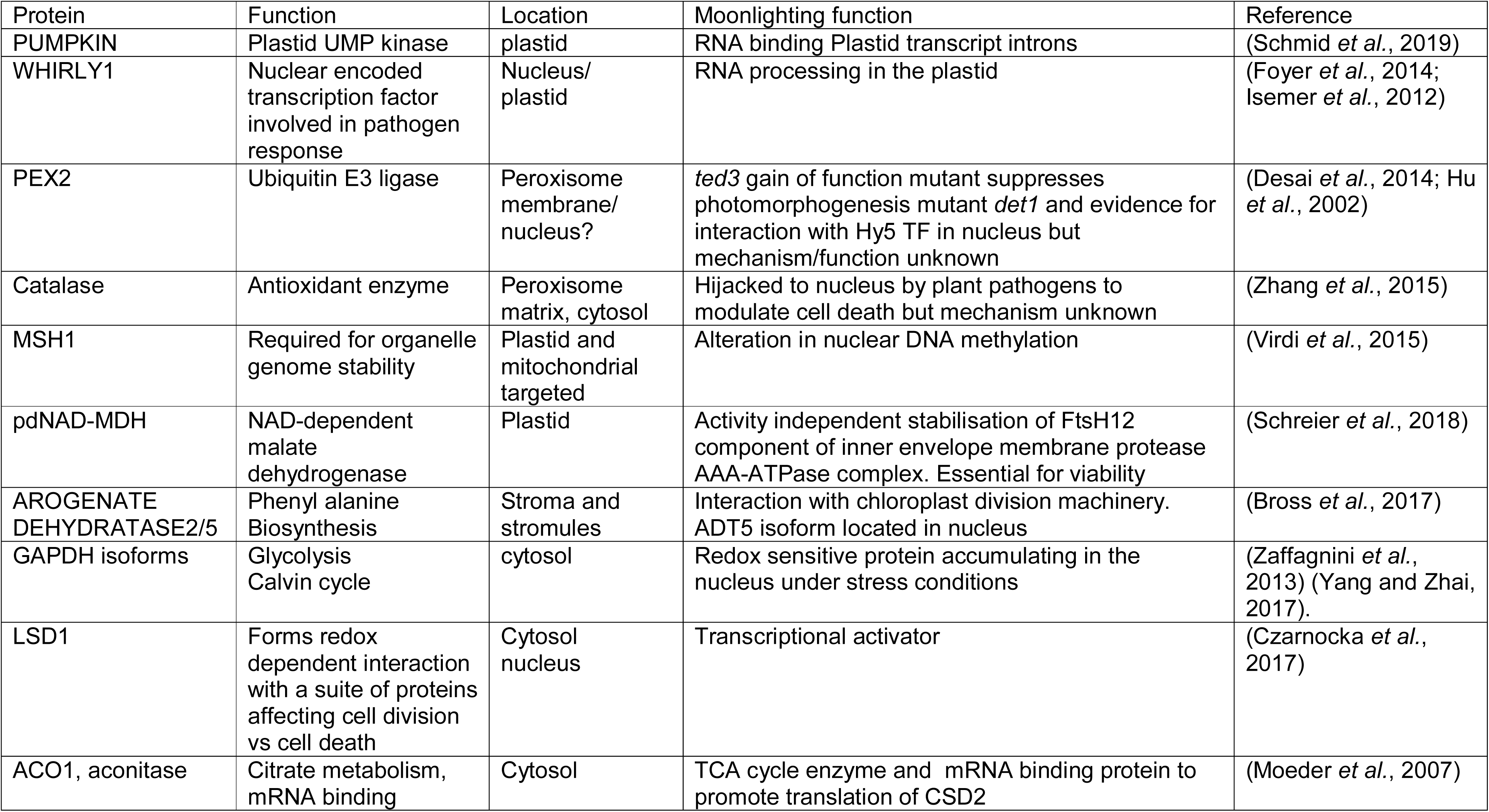
Moonlighting proteins in plants

| Protein | Function | Location | Moonlighting function | Reference |
| --- | --- | --- | --- | --- |
| PUMPKIN | Plastid UMP kinase | plastid | RNA binding Plastid transcript introns | (Schmid *et al.*, 2019) |
| WHIRLY1 | Nuclear encoded transcription factor involved in pathogen response | Nucleus/<br>plastid | RNA processing in the plastid | (Foyer *et al.*, 2014; Isemer *et al.*, 2012) |
| PEX2 | Ubiquitin E3 ligase | Peroxisome membrane/<br>nucleus? | *ted3* gain of function mutant suppresses photomorphogenesis mutant *det1* and evidence for interaction with Hy5 TF in nucleus but mechanism/function unknown | (Desai *et al.*, 2014; Hu *et al.*, 2002) |
| Catalase | Antioxidant enzyme | Peroxisome matrix, cytosol | Hijacked to nucleus by plant pathogens to modulate cell death but mechanism unknown | (Zhang *et al.*, 2015) |
| MSH1 | Required for organelle genome stability | Plastid and mitochondrial targeted | Alteration in nuclear DNA methylation | (Virdi *et al.*, 2015) |
| pdNAD-MDH | NAD-dependent malate dehydrogenase | Plastid | Activity independent stabilisation of FtsH12 component of inner envelope membrane protease AAA-ATPase complex. Essential for viability | (Schreier *et al.*, 2018) |
| AROGENATE DEHYDRATASE2/5 | Phenyl alanine Biosynthesis | Stroma and stromules | Interaction with chloroplast division machinery. ADT5 isoform located in nucleus | (Bross *et al.*, 2017) |
| GAPDH isoforms | Glycolysis Calvin cycle | cytosol | Redox sensitive protein accumulating in the nucleus under stress conditions | (Zaffagnini *et al.*, 2013) (Yang and Zhai, 2017). |
| LSD1 | Forms redox dependent interaction with a suite of proteins affecting cell division vs cell death | Cytosol nucleus | Transcriptional activator | (Czarnocka *et al.*, 2017) |
| ACO1, aconitase | Citrate metabolism, mRNA binding | Cytosol | TCA cycle enzyme and mRNA binding protein to promote translation of CSD2 | (Moeder *et al.*, 2007) |

## Abbreviations

ER: endoplasmic reticulum
CAT: catalase
GAPDH: glyceraldehyde 3- phosphate dehydrogenase
NPR1: NON-EXPRESSOR OF PATHOGENESIS-RELATED GENES 1;
PR: pathogenesis-related
PRX: peroxiredoxins
PTM: post-translational modifications
ROS: reactive oxygen species
RNS: reactive nitrogen species, SA: salicylic acid
SAR: systemic acquired resistance
SOD: Superoxide Dismutase
TRX: thioredoxin
UPS: Ubiquitin-proteasome system
WHY1: WHIRLY1

## Author contributions

CH. Foyer and A Baker proposed the original idea and wrote the first draft of the manuscript text. All authors polished and contributed to the final manuscript text. A Baker produced figures 1 and 2, and table 1.J. Schippers produced Figure 3 and harmonised the style of all the figures. M Wright produced Box I.

## Acknowledgements

The authors gratefully acknowledge Prof Patricia Conklin State University of New York College at Cortland for the image in Figure 4. CHF thanks BBSRC (UK) for financial support (BB/N004914/1).

## Box I - key developments to help understand reversible oxidative modifications in plants

### 1. Genetically-encoded protein-based tools to trap sulfenylated proteins *in situ*

*S*-Sulfenylation (protein-SOH) is a reversible oxidative PTM that acts as regulatory switch in signal transduction pathways. However the global “sulfenome” is particularly challenging to detect as this PTM is transient, unstable, and prone to over-oxidation even during cell lysis. Recently a genetically-encoded tool to capture *S*-sulfenylated proteins was developed (Waszczak *et al.*, 2014). The cysteine-rich domain of the yeast transcription factor YAP1 forms disulfides with *S*-sulfenic acid modifications on its cognate signalling protein; fusion of this domain with an affinity tag creates a tool to capture and enrich *S*-sulfenylated proteins *in vivo* (Figure 5, **A**). YAP1 can be expressed in cells, with control cells expressing a catalytically inactive version (YAP1A), and, following cell lysis, downstream affinity purification used to identify disulfide linked proteins. The authors detected ∼100 sulfenylated proteins in Arabidopsis cell suspensions exposed to H_2_O_2_ oxidative stress (Waszczak *et al.*, 2014).

### 2. Small molecule-based probes to detect the sulfenome

A complementary approach exploits the chemoselective reaction of small molecules based on dimedone with sulfenic acid. Whilst YAP1C recognition of sulfenic acids is dependent on protein-protein interactions, a small molecule is in principle more general and able to access more sulfenylation sites. The Carroll group have pioneered the use of DYn-2, a dimedone probe that is small yet ready appended to affinity tags such as biotin by click chemistry for enrichment of sulfenylated proteins (Figure 5, **B**) (Paulsen *et al.*, 2011). Akter et al. applied DYn-2 is Arabidopsis cultures (Akter *et al.*, 2015), identifying 226 sulfenylation events in response to oxidative stress, and, more recently, in plants (Akter *et al.*, 2017).

